# Nucleosome-mediated conformational switches in micro-eccDNAs

**DOI:** 10.1101/2025.06.14.659616

**Authors:** Sarah Harris, Victor Velasco Berrelleza, William Sandel, Thana Sutthibutpong, Vicki Parikh, Euan Ashley, Andrew Fire, Stephen D. Levene, Thomas C Bishop, Massa Shoura

## Abstract

Extrachromosomal-circular DNA (eccDNA) are circular-DNA elements that are implicated in cellular processes such as genomic diversification and instability, along with other genomic mechanisms. However, their structure and chromatinization state is poorly understood. Here we identified a 358-bp circular micro-eccDNA molecule derived from the human Titin gene, which was selected for this study based on its size and sequence characteristics, and used atomistic molecular dynamics simulations to study its interaction with nucleosomes and corresponding topological state. We show the presence or absence of bound nucleosomes provides a topological switch between intact and denatured DNA.

## GRAPHICAL ABSTRACT

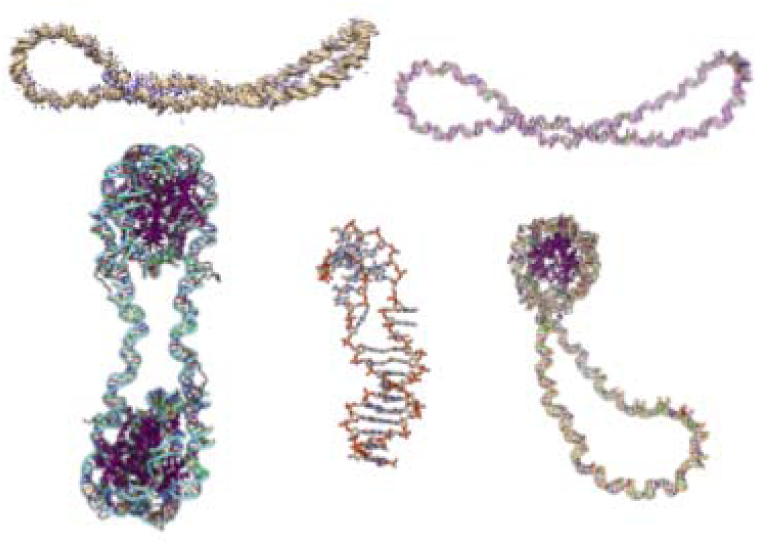

## INTRODUCTION

Extrachromosomal-circular DNA (eccDNA) comprises a diverse class of endogenous circular-DNA elements that are implicated in genomic diversification and instability, programmed genetic rearrangements, gene amplification, structural variation, cellular senescence, and other processes. These circular molecules, which we collectively denote the “Circulome”, derive from segments of the canonical linear genome and span an extremely wide range of molecular sizes from ≤150 bp to > 1·10^6^ bp. While their sequence composition reflects the genome from which they are derived, pathological state, and developmental stage, eccDNAs can acquire additional complexity, including unique mutations. Global mapping of eccDNA- sequence data by high-throughput sequencing shows that eccDNAs originate from discrete and reproducible locations in both the wild-type C. elegans and both normal and diseased human tissues(1) (2). However, the repertoire of the circulome is highly dynamic, varying by cell type, disease state, and selective pressures. Notably, eccDNAs harboring genetic and regulatory elements have been implicated in oncogene amplification in certain cancers (3) (4–6) (7) (8) (9) (10) (11) (12) (13). Although our understanding of detailed mechanisms responsible for eccDNA biogenesis in different cellular backgrounds remains fragmentary, models for multiple subsets of eccDNAs have been proposed (3) (14) (15) (16). Depending on mechanistic details, formation of eccDNA can be associated with gene-amplification and/or deletion events on the linear DNA. This highlights the biological consequences of eccDNA formation beyond the direct functional roles these circular molecules can play in the cell.

The transcription capabilities (and transcription levels) of eccDNA-borne gene segments remain controversial, particularly in the case of ultra-small eccDNAs (a.k.a microDNAs;100 bp-500 bp). These molecules do not contain full genes, but have been proposed to encode microRNAs that post-transcriptionally regulate cognate loci on the linear genome (17) through “eccDNA-mediated gene silencing,” even in cases where eccDNAs do not harbour canonical promoter sequences. Moreover, the identification of micro-peptides as agents of biological activity also raises the intriguing possibility that micro-eccDNAs may be capable of producing such peptides, particularly as the transcripts need not contain the expected AUG start codon (18) (19). The potential for micro-eccDNAs to produce biologically active small RNAs and micro-peptides depends upon the transcription and the expression levels of the circular template. We hypothesize that the most favorable state for these ultra-small eccDNAs to support “promoter-less” transcription is through the formation of denatured- DNA bubbles containing single stranded DNA regions in unchromatinized segments of the micro-eccDNA, thereby providing the transcription machinery with access to the genetic code.

The question of whether or how eccDNAs in cells are organized as chromatin and the dynamics of these structures remains open. Based on ATAC-seq data, a subset of eccDNAs (multi-genic, hence larger in size) in glioblastoma cell lines appear to harbour nucleosomes, albeit at an apparent reduced density compared with their cognate loci on the linear genome.(3) At the opposite extreme of the circulome’s molecular-size distribution, micro- eccDNAs become comparable in size to that of a mononucleosomal DNA. The presence of nucleosomes in populations of small micro-eccDNAs has not yet been assessed experimentally because the methods used thus far to analyse micro-eccDNA require protein-free DNA. However, nucleosome-reconstitution experiments with DNA have shown that it is possible to assemble nucleosomes on small circular-DNA templates, comparable in size to that investigated in the present study (20) (21).

To investigate whether micro-eccDNAs in this size range can exist in a chromatinized state, we focus on questions of nucleosome occupancy for a promoter-less 358-bp micro-eccDNA element we have discovered, which maps to an exon from the human Titin (TTN) gene. Using molecular dynamics (MD) simulations, we address: (i.) whether a micro-eccDNA molecule in this size range can accommodate up to two nucleosomes; (ii.) effects of nucleosome loss on duplex stability; (iii.) how DNA transcription/accessibility can in turn be modulated by micro-eccDNA topology and chromatin structure. MD simulations of DNA minicircles have been used in conjunction with cryo-EM (22) and AFM (23) experiments to show in atomic detail how the structure and dynamics of the DNA is affected by DNA supercoiling and topology. Here we use MD simulations to compare structures of the di- and mononucleosome-containing 358-bp micro-eccDNA with the naked micro-eccDNA, subject to the assumption that the initial dinucleosome structure is topologically relaxed. Whereas the relaxed dinucleosome-bearing micro-eccDNA remained double-stranded, we observed formation of denaturation bubbles in this molecule subjected to removal of one or both nucleosomes. Thus, we conclude that for these micro-eccDNAs to support promoter-less transcription of small RNAs and potentially contribute to eccDNA-mediated gene silencing, they must exist in a partially or fully dechromatinized state.

## MATERIAL AND METHODS

### Experimental Details

Dermal fibroblasts were isolated from participating patients and passaged three times before lentiviral infection with OCT4, SOX2, KLF4, and c-MYC. Colonies with iPSC morphology were selected, isolated and maintained on Matrigel-coated plates (BD Biosciences) for maintenance with mTESR-1 growth medium (StemCell Technologies) as previously described (24). After at least 20 passages, iPSCs were split 1:15 into 24 well plates. Four days following plating, when the cells reached 90% confluency, differentiation was induced with 0.4 µM ChiR (Sellek Chemicals) in RPMI with L-glutamine and glucose (Corning), 5% B27 supplement without insulin (Fisher Scientific) for two days followed by 2 days treatment with 0.5 µM IWR (Sigma) in the same media. Cells were maintained in RPMI with L-glutamine and glucose (Corning) with 5% B27 supplement (Fisher Scientific) for 12 days and raised in RPMI without glucose, with 6mM lactate and 5% B27 supplement (Fisher Scientific), with media changed every third day. Genomic DNA was extracted from each cell line at the iPSC stage, and at 20, 30 and 40 days after the start of differentiation. At each time point, adherent cells were lifted with Versene (Gibco), pelleted and DNA was extracted using Blood and Cell Culture DNA Mini Kit (Qiagen). eccDNA was isolated from genomic DNA using prolonged and progressive ExoV treatment, following our RCA-free Circulome-Seq protocol(1). Post enrichment, libraries were then prepared from the enriched eccDNAs using an optimized attenuated Tn5 tagmentation approach (Nextera XT, Illumina) designed specifically for singly tagmented circles, as described in (1) without any prior amplification. Paired-end sequencing of singly tagmented circles enabled the complete assembly of eccDNA sequences for each uniquely cut molecule. Among these, we identified a 358-bp circular DNA molecule derived from the Titin gene, which was selected for this study based on its size and sequence characteristics.

### Model Building for MD simulations

In-silico models of the dinucleosome structure based on the 358 bp Titin micro-eccDNA sequence were built using the Interactive Chromatin Modeling Webserver (25) containing two nucleosomes each with a 147-bp core and 16 bp of linker-DNA arms flanking the core. The linker DNA was constructed using relaxed B-form DNA with adjustments to ensure that the DNA could be connected, and the nucleosomal region was constructed using the structure of 1KX5 including tails (26), with the sequence replaced by that of the Titin micro- eccDNA. Nucleosome tails were modified by methylation of K9 and K27 on histone H3. Simulations in explicit solvent were carried out in the presence of either monovalent 1:1 salt (300 mM NaCl) or iso-Coulombic divalent-salt conditions (150 mM CaCl_2_) as described below. MD simulations were performed on NVIDIA V100 GPUs, which provided 4.9 ns per day for the largest naked-DNA simulations containing ∼2.7 million atoms.

### Multi-scale MD Simulation Equilibration Scheme

All atomistic MD simulations employed a combination of implicit and explicitly solvated calculations using the AMBER 20 suite of programs (27) (28). Implicit-solvent simulations using the AMBER GB/SA methodology effectively accelerate MD timescales due to the absence of solvent damping (so long as the thermostat is only weakly coupled), meaning that only a few nanoseconds of implicit MD can explore conformational changes that would require ∼100 ns to observe in explicit water (22). However, simulations in implicit solvent are less structurally robust than when the system is fully solvated and provide only an approximate representation of ion-DNA interactions.

Consequently, we employ a multiscale simulation scheme whereby micro-eccDNAs are first equilibrated in implicit solvent in the presence of hydrogen-bond restraints over timescales of 1 −2 ns, followed by a full explicitly solvated MD run for > 100 ns (see Table 1) with all restraints removed. Structures were selected from the implicitly solvated MD trajectories after sufficient time that the twist-writhe partition had equilibrated (see Supplementary Figure 1), but prior to any disruption in base stacking, which we observe over longer timescales. To ensure that the quality of our explicitly solvated structures is not compromised by the implicit solvent equilibration, we discard solvated simulations that exhibit defects over a timescale shorter than 10 ns. This assumes that defects that arise a sufficiently long time after equilibration are not artifacts of the equilibration process. For example, we discarded four naked micro-eccDNA trajectories from our analysis based on this criterion.

### Dinucleosome-microDNA Simulation Equilibration

The dinucleosome-DNA complex was first neutralised with Ca^2+^ ions, a periodic box of 10 Å of TIP3P waters was added, and the number of excess ions was calculated by comparing 150 mM Ca^2+^ to the 55.5 moles/liter density of water molecules. Equilibration of the solvent and DNA prior to all production runs used a standard multistage protocol (29). To compare the behaviour of micro-eccDNA in divalent and monovalent ions, while maintaining the same electrostatic environment, after 100 ns MD each divalent Ca^2+^ cation was replaced by a monovalent Na^+^ ion in the same location; and additional Na^+^ cations were added in random positions, thereby maintaining iso-Coulombic conditions. MD was then performed for a subsequent 100 ns.

### Mononucleosome-microDNA Simulation Equilibration

A single nucleosome was deleted from the dinucleosome-microDNA complex and equilibrated in implicit solvent for 2 ns in the presence of hydrogen-bond restraints. Solvent and 150 mM CaCl_2_ were added, and a 100 ns unrestrained production run was performed. The stability criterion of maintaining a 10 ns defect-free post-equilibration solvated structure was difficult to achieve, in spite of four different attempts with starting structures sampled from the implicit MD.

### Naked microDNA Simulation Equilibration

Both nucleosomes were deleted from the dinucleosome complex and equilibrated in implicit solvent for 1.5 ns in the presence of hydrogen-bond restraints, before solvent and 150 mM CaCl_2_ or 300 mM NaCl were added. This was followed by a 175-ns production run. Longer unrestrained MD runs were required in the absence of nucleosomes because the changes in writhe that occur between monovalent- and divalent-cation conditions involve slow structural relaxation at the crossing points of plectonemes. An additional 150 ns simulation in 150 mM CaCl_2_ was performed starting from the DNA structure obtained after 2 ns of implicitly solvated MD. At the end of this simulation, the divalent Ca^2+^ ions were replaced with Na^+^ and additional Na^+^ ions were added in random positions as described previously, and unrestrained MD was performed for a subsequent 150 ns in 300 mM NaCl.

### Simulation Analysis Methods

The twist (*Tw_i_*) at each base-pair *i* and writhe of the eccDNA were calculated using the WrLine program (30). We define the total twist (*Tw*) in DNA turns as the sum of base-pair twist values:

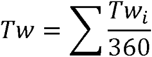

The linking number difference □*Lk* and superhelix density *σ* were calculated using:

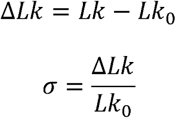

where *Lk* = *Tw + Wr*, and *Lk_0_* is assumed to be the number of turns of a relaxed B-DNA structure of the same length: *Lk_0_* = 358/10.5.

Ion-density analysis was performed using CCPTRAJ in AMBER20. Ion-density maps were calculated from the last 75 ns of each production run. Hydrogen bonding (defects) analysis was performed by calculating the average distances between two heavy atoms among complementary base paired atoms using MDAnalysis (31) (32) in Python (see Figure S5).

## RESULTS

Following isolation and sequencing, the 358-bp Titin circular micro-eccDNA characterized experimentally was constructed in silico, and investigated in the presence of zero (i.e., naked), one, and two nucleosomes using atomistic MD simulations in explicit water. When building the computational models, we make the assumption that when the micro-eccDNA is excised from the genome, it contains the maximum number of nucleosomes possible, which in this case is two. We also assume that this maximally chrominatised microDNA circle was in a relaxed state when excised. Analysis of the resulting simulated micro-eccDNA structures with the WrLine program shows that the 358-bp Titin circular DNAs constructed from a combination of nucleosome core particles and relaxed linker-region DNA has □Lk ∼ −2.15 and σ ∼-0.063 (see Supplementary Figure S2). These levels of negative supercoiling are close to those observed in bacteria in vitro when DNA-binding proteins are removed (33) (34), and have been experimentally observed to support the formation of denatured regions in DNA minicircles of similar sizes (35).

Table 1 provides details of the explicitly atomistic MD simulations performed for the mono and dinucleosome and naked micro-eccDNAs. We simulated the dinucleosome-microDNA for 100 ns and naked micro-eccDNAs for 175 ns, and compared behaviour in divalent ions (150 mM CaCl_2_) with monovalent ions (300 mM NaCl) in both cases. For the naked DNA, additional simulations were performed to check the sensitivity of the results to the initial structures because significant differences in the twist-writhe partitioning between mono- and divalent cations was observed. However, only one 100-ns trajectory in divalent ions for the mononucleosomal micro-eccDNA is reported because significant disruption to the complementary base-pairing interactions occurred from the beginning of the MD simulation.

### Twist-writhe partitioning in nucleosome-containing and naked micro-eccDNA simulations

The time evolution of twist-writhe partitioning within the micro-eccDNAs in explicit solvent MD is shown in Figure 1. Figure 2 shows representative structures comparing the dinucleosome (Figure 2a) with the mononucleosome (Figure 2b) and the naked DNA (Figure 2c,d). When the nucleosomes are removed, the resulting electrostatic repulsion causes strikingly large rearrangements of the micro-eccDNA structure into conformations that are significantly less compacted by writhing. Supplementary Movies 1 and 2 show the relaxation of the DNA observed in the implicit solvent MD simulations when one and two nucleosomes are removed respectively. The writhe is reduced from approximately −2 for the relaxed dinucleosome-eccDNA to −1.5 when one nucleosome is removed. When both nucleosomes are removed, the writhe of the resulting naked micro-eccDNA relaxes to between −1.5 and - 0.5, depending upon the counterion environment (see section 3.2) and the degree of denaturation in the DNA (see section 3.3).

**Figure 1:**
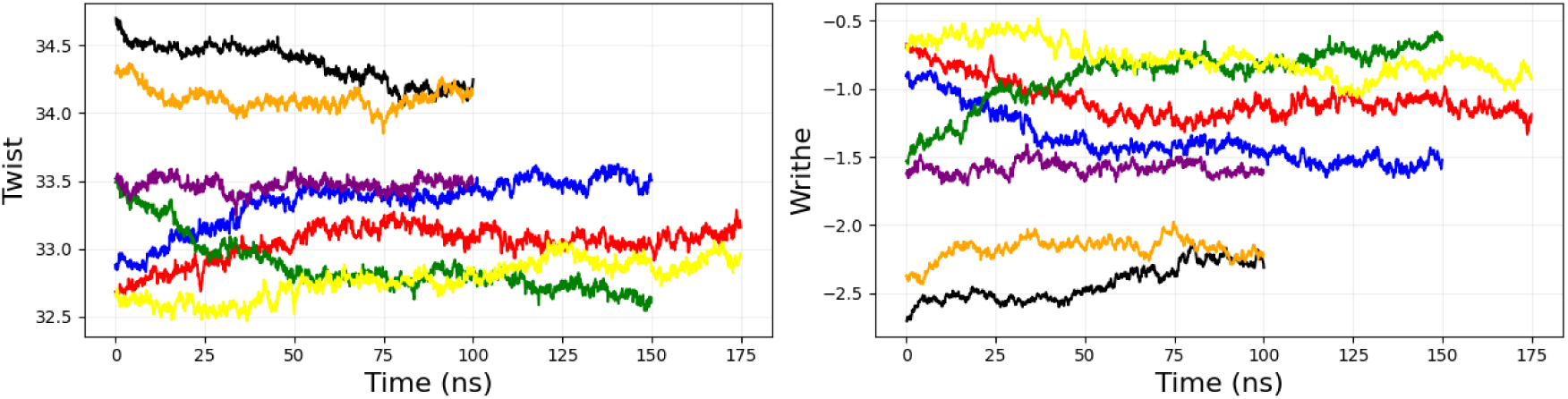
Time dependence of the twist and writhe in explicitly solvated MD simulations showing dinucleosome in CaCl_2_(black) and NaCl (orange); mononucleosome in CaCl_2_ (purple); and naked eccDNA in CaCl_2_ (blue/red) and in NaCl (yellow/green).

**Figure 2:**
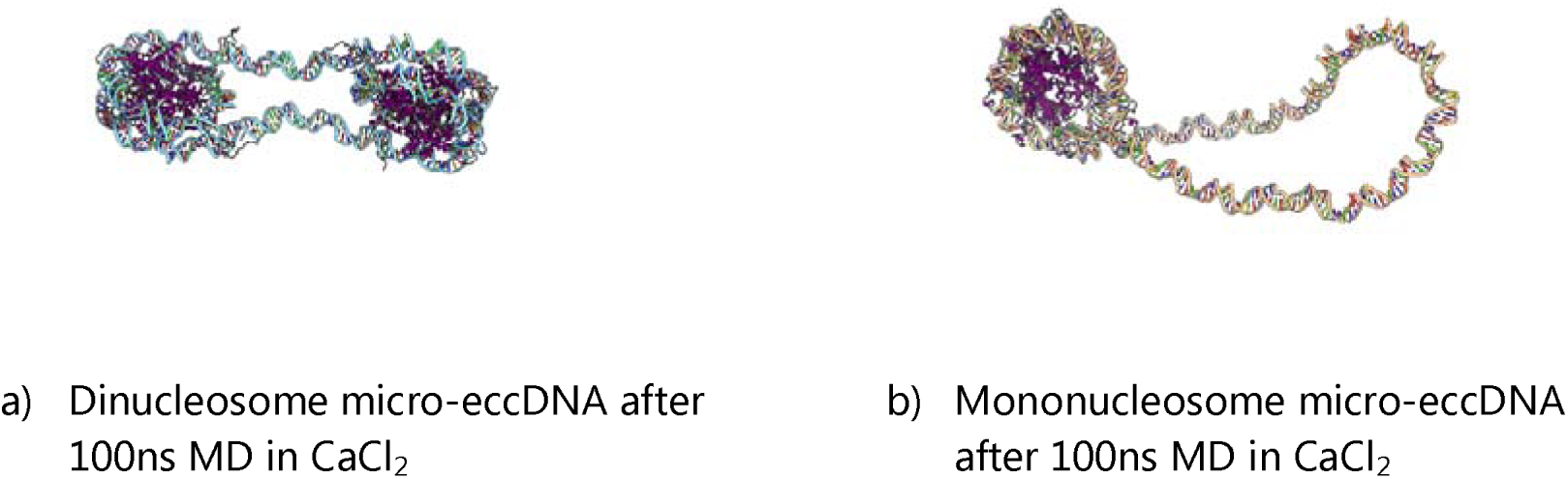
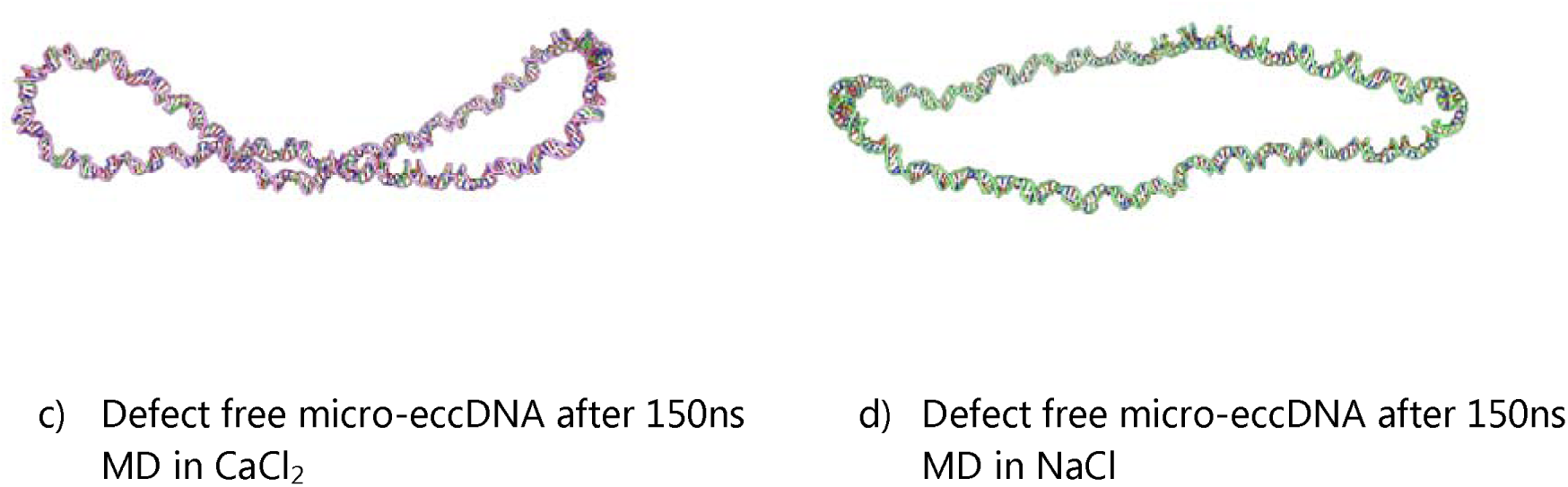
Representative structures of micro-eccDNA in the presence and absence of nucleosomes after production MD. The structures are shown to scale, so that the degree of compaction of the circles when nucleosomes are bound is clear. Defect-free micro-eccDNAs are illustrated here because these have the greatest difference in writhe between mono and divalent cations.

### Micro-eccDNA-conformation-dependent ion distributions in MD simulations

Figure 2 compares micro-eccDNA structures containing one (Fig 2b) and two (Fig 2a) nucleosomes after 100 ns of simulation in divalent (Ca^2+^) ions with naked micro-eccDNA in monovalent (Na^+^) (Fig 2c) and divalent ions (Fig 2d). While no significant structural differences between mono and divalent ions were observed for the dinucleosome, the naked micro-eccDNAs were able to relax to more negative writhe values in divalent compared to monovalent ions. In divalent ions, the two strands are annealed at the crossing point through a Ca^2+^ ion bridge (see ion density map in Fig. 3a and supplementary Fig. S3), enabling a more writhed structure (Figure 1 blue/red) than for monovalent Na^+^ ions (Figure 3b and Figure 1 green/yellow). Repeating these simulations from independent starting structures and recording where non-neighbouring DNA strands are 10 Å apart or less shows that the location of the close crossing is variable (see Supplementary Figure 4), so in this micro- eccDNA the positions of the plectoneme tips are not uniquely specified by the sequence.

**Figure 3:**
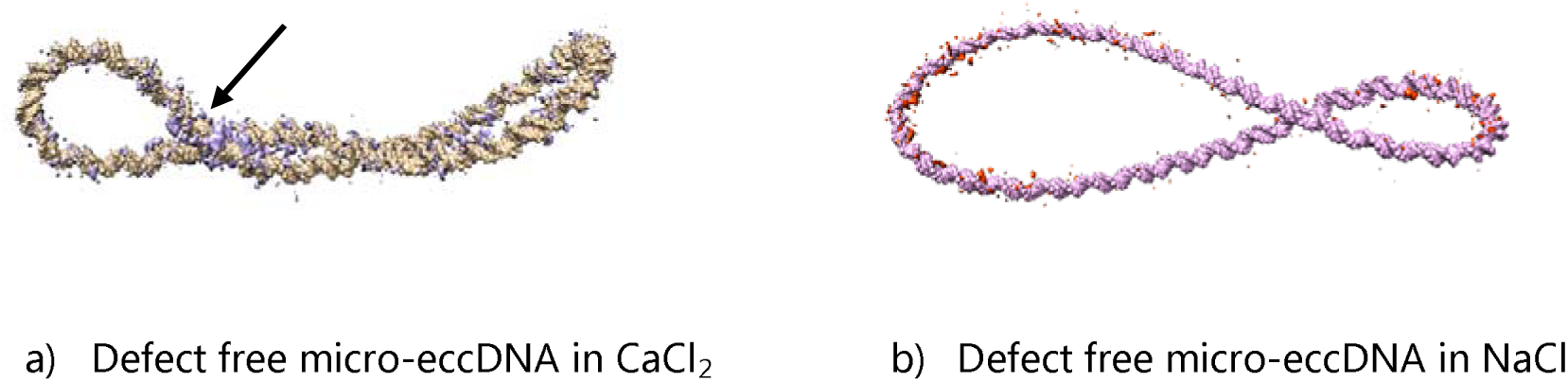
Ion density maps for a) CaCl_2_ solution (density of Ca^2+^ shown in violet) and b) NaCl (Na^+^ density shown in red) superimposed on the time-averaged DNA structures. The region of high divalent-ion density observed at the crossing point is indicated with an arrow.

### Base-pair stability in nucleosome-containing and naked micro-eccDNA simulations

Figure 4 shows the base-pair distances between complementary base pairs as a function of time during each of the MD trajectories. Large excursions of inter-base-pair distances, as shown in yellow, are signatures of transient local defects in helical structure and base-pair separation or denaturation.

**Figure 4:**
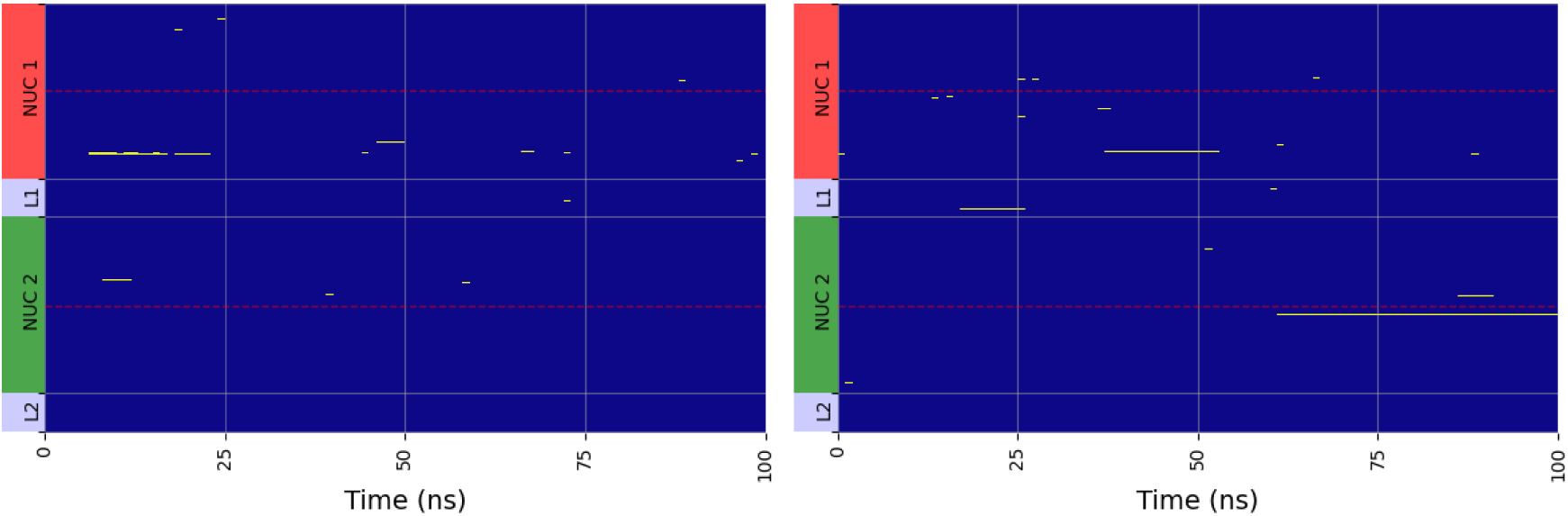
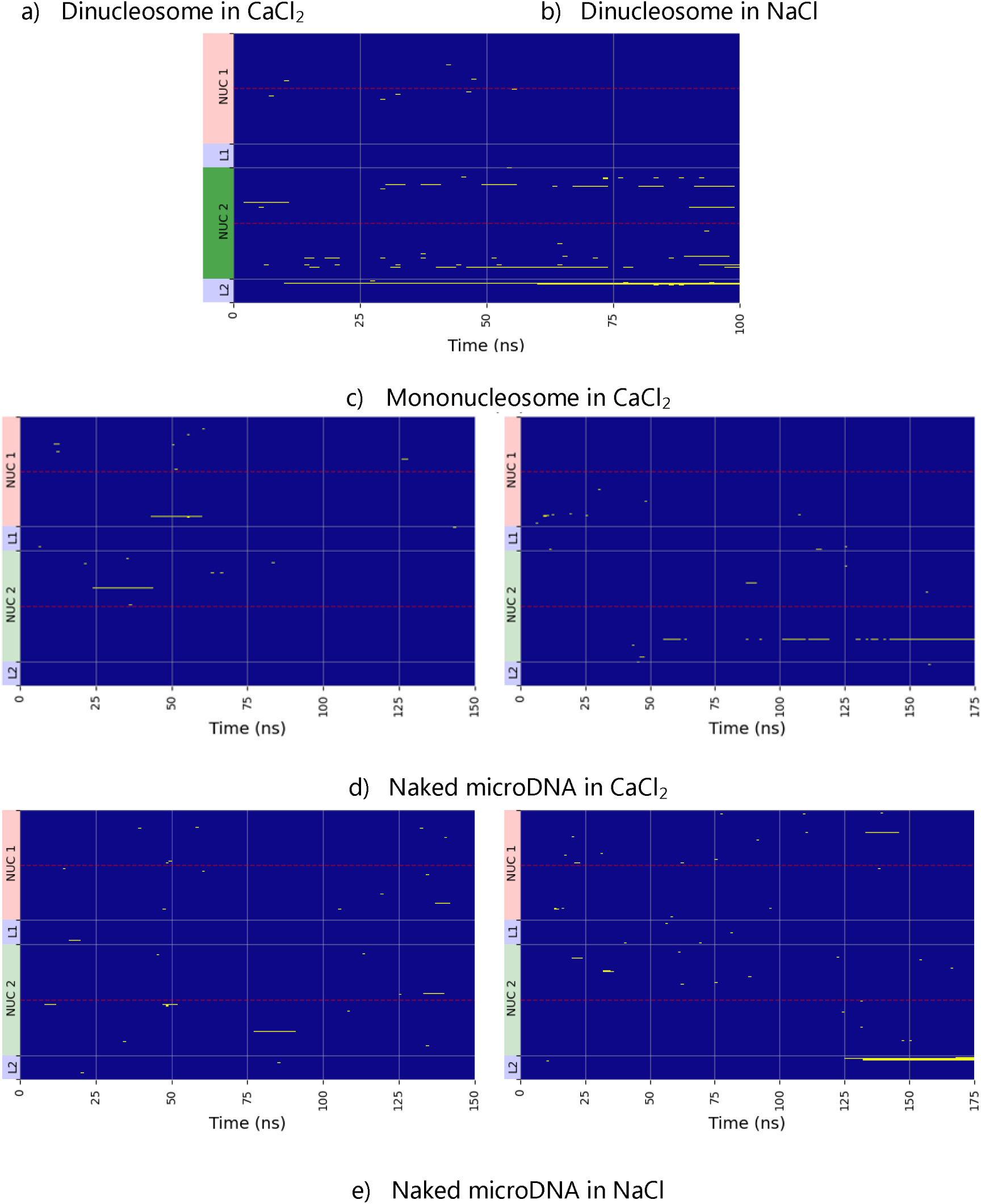
Time dependence of the distances between complementary base pairs during solvated MD. Yellow indicates broken base pairs (distances between heavy atoms > 4Å). For reference, the nucleosome positions are indicated in red/pink or dark/light green (occupied/unoccupied); red and white horizontal lines indicate nucleosome centres and edges respectively.

In the dinucleosome-containing micro-eccDNAs, the double helix remains predominantly intact (see Fig 4). However, in the mononucleosome-containing micro-eccDNA there is significant denaturation. Broken base pairs occurred after less than 10 ns, implying that these micro-eccDNAs are intrinsically unstable, presumably because the empty domain of the eccDNA is too small to relieve supercoiling stress by writhing. Inspection of the structures indicates that this occurs predominantly at the highly bent region where the DNA exits from the bound nucleosomal region (see Figure 5). In naked micro-eccDNAs, a combination of transient opening and denaturation events are observed. In the naked micro-eccDNAs, prominent denatured regions occur close to plectoneme apices (36) (37), where strong bends contribute to base-pair disruption (see Figure 5).

**Figure 5:**
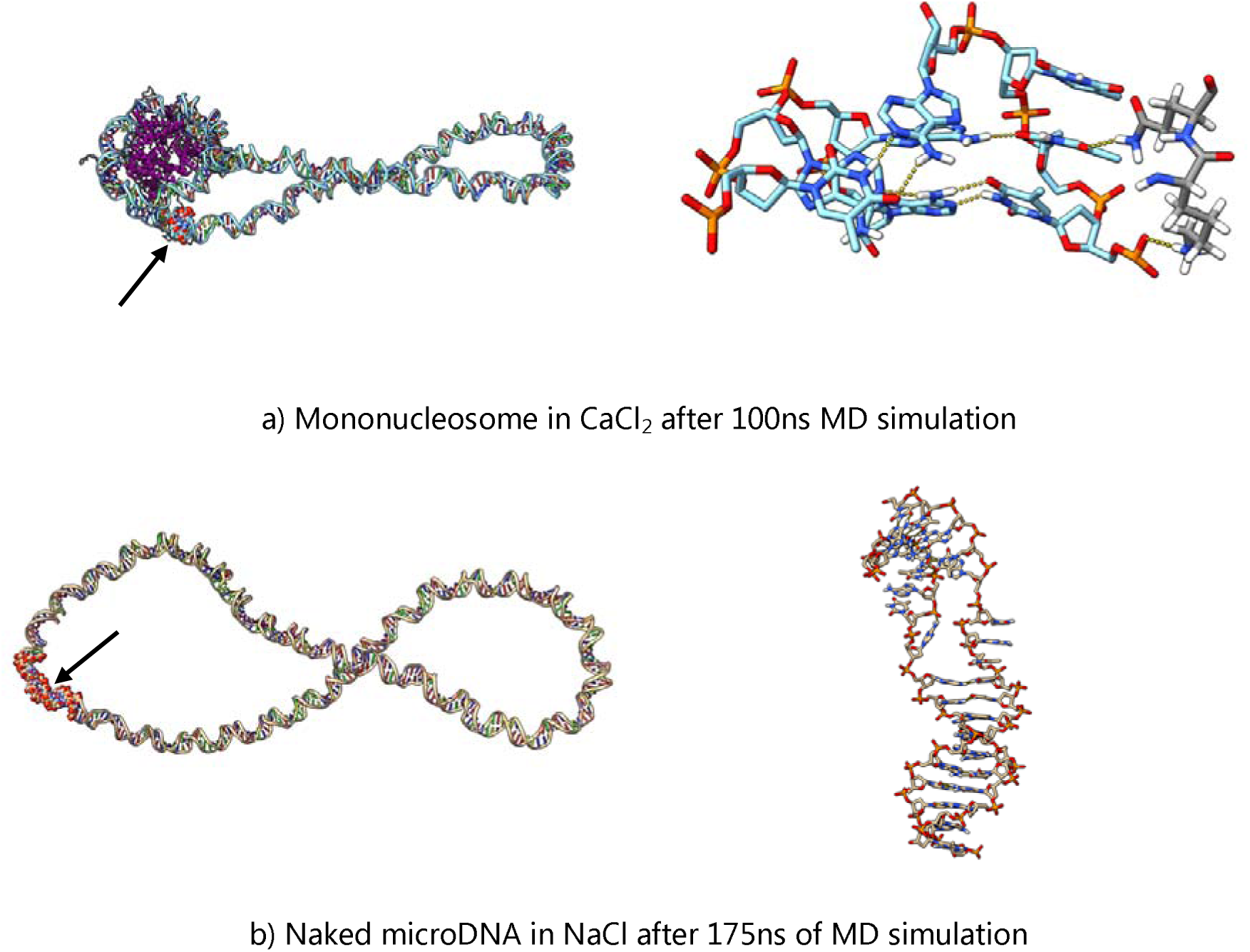
Examples of denaturation between complementary base pairs observed in the final structures sampled in the MD trajectories, showing the full structures on the left and close-up views of the denatured region on the right. The denatured regions are shown as spheres and indicated by an arrow on the left, hydrogen bonds are shown in yellow in the zoomed-in structures on the right. In the mononucleosome the DNA forms hydrogen bonds with lysine and glutamine residues within the nucleosome tails, shown in grey.

Figure 6 shows the timescale of opening events for AT and GC base pairs. Of all 1313 opening events, 12 persisted beyond the timescale of the simulation, with 11 occurring in AT base pairs. All of the transient opening events last for a maximum of around 20 ns. Apart from fraying at the duplex ends, breathing events are almost entirely absent from long (µs) MD simulations of linear DNA sequences (38), and must therefore be a result of negative supercoiling.

**Figure 6:**
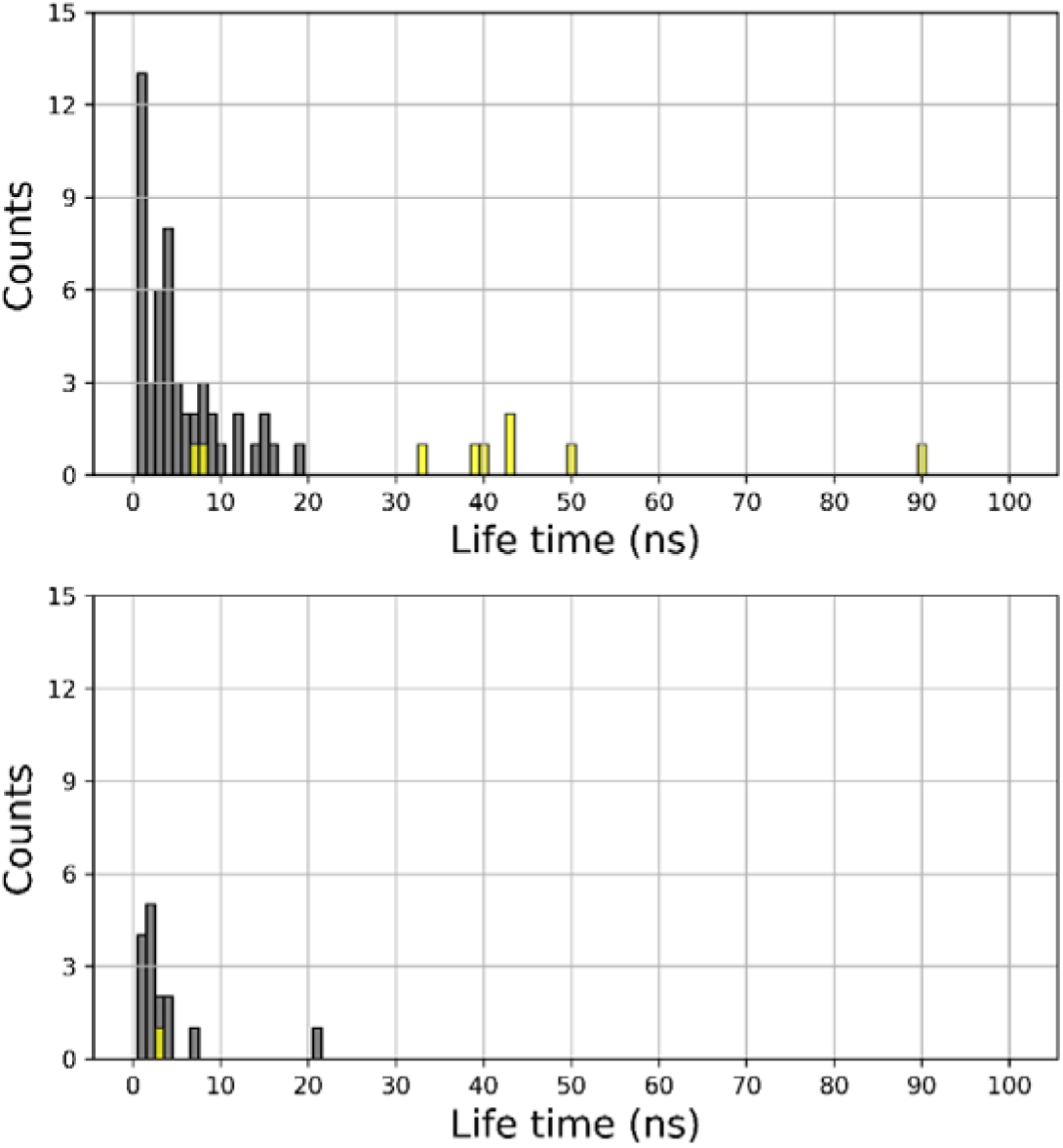
Timescale of opening events for AT (top) and GC (bottom) base pairs. Grey bars indicate base pairs opening events that repair over the timescale of the simulation, whereas yellow bars show events that persist beyond the length of the MD trajectory. Note that shorter lived opening events that are not observed to reanneal occur towards the ends of the MD runs.

## DISCUSSION

Extrachromosomal-circular DNA (eccDNA) is a component of genomes consisting of circular molecules that range widely in size, from ∼ 100 bp to > 1 million bp (3). In normal cells, eccDNA populations can contain many small DNA rings and the recovery of these small DNA species in the bottom band of cesium chloride/ethidium bromide density gradients indicates that these molecules are covalently closed (1). Whether such ultra-small DNA circles are organized into minicircle chromatin structures in vivo remains an important and open question. Moreover, it has been proposed that micro-eccDNAs can act as templates for transcription without the need for canonical promoters. Conversion of stored superhelical free energy in the micro-eccDNA into localized DNA unwinding is a potential mechanism for regulating micro-eccDNA transcription in the absence of a canonical promoter sequence because this results in the formation of a denatured single-stranded region of DNA that can provide access to the transcription machinery. The work described here is thus an initial effort to computationally address these issues from the perspective of nucleosomal DNA structure and dynamics, using an experimentally observed micro-eccDNA sequence.

Micro-eccDNAs containing less than 400 bp can, in principle, accommodate up to two nucleosomes. Although nucleosomes were placed equidistantly along the circular-DNA contour, the register of the DNA was assigned arbitrarily, and presumably is not able to equilibrate over the timescale of the simulations due to the strong electrostatic interactions between the DNA and the nucleosome surfaces. We therefore suggest that future calculations performed at a coarse-grained level could be used to provide alternative conformations, which could then be compared using more computationally intensive atomistic simulations so that the potential importance of register can be assessed.

Starting from the relaxed dinucleosomal micro-eccDNA, we investigated effects of instantaneously removing one or both histone octamers. Removing all histones from a relaxed 358-bp dinucleosomal circle results in a DNA ring with negative supercoiling comparable to that observed in protein-free bacterial DNA (33) (34). We find that dinucleosomal micro-eccDNA models retain almost all hydrogen bonding interactions between complementary base pairs over the course of our MD simulations, whereas mononucleosome-containing circles were highly prone to denaturation. The residual negative supercoiling resulting from these perturbations can approach the superhelix density of bacterial plasmids in vitro and, in experiments using protein-free DNA minicircles, were sufficient to induce increased base breathing and locally denatured duplex regions. Consistent with these experimental observations we observed stable localized unwinding in dinucleosomal micro-eccDNAs that underwent instantaneous nucleosome eviction. Notably the unwound regions were not reproducibly associated with particular sequence elements, although more extensive sampling and larger datasets could point to specific trends.

Partitioning between twist and writhe is modulated by environmental conditions, with more highly writhed and compact minicircle structures being favored by divalent ions when compared to monovalent ions at comparable ionic strength. To our knowledge, this behavior has not been addressed in previous MD simulations of micro-eccDNAs, although previous studies of idealised geometries have indicated that left handed crossovers in plectonemes should be stabilized by divalent ions (39,40). Cryo-TEM experiments on plasmid-sized circular DNAs, which are about an order of magnitude larger than the minicircles investigated here, showed that divalent ions promote superhelical-DNA compaction, consistent with the divalent-ion effects that we observe in simulated minicircles (41). Subsequent coarse-grained Brownian-dynamics simulations from the same group provided a basis for understanding specific effects of ion valence on conformational distributions of plasmid-sized superhelices (42). Thus, the observed compaction of the nucleosome-free micro-eccDNA as a function of ion composition in our simulations qualitatively recapitulates and generalizes behavior observed in larger DNA molecules.

Given these results, we can evaluate the hypothesis regarding the capacity of ultra-small eccDNAs for promoter-less transcription and potential biological roles, for example, gene silencing via small RNAs, or production of micro-peptides. Our findings suggest that for these micro-eccDNAs to be transcribed without a canonical promoter—potentially through the formation of a DNA bubble, they must adopt to first order a specific topologically dechromatinized state or act as a topological switch by dynamically transitioning between a chromatinized and a less chromatinized state. There remain many open questions concerning the chromatin state of micro-eccDNAs, not least whether nucleosomal forms of these circles exist in the cell. Although the presence of nucleosomes in large eccDNAs has been directly probed, technical challenges have for the moment limited our ability to assay for smaller nucleosomal micro-eccDNAs. Moreover, if nucleosomal micro-eccDNAs exist, what is their epigenetic state? In our simulations K9 and K27 of histone H3 were methylated; however, the space of possible histone modifications is huge and in the case of eccDNA, effectively unexplored. Histone modifications have profound effects on nucleosome stability and inter-nucleosomal interactions (43). DNA-base modifications are also possible in the case of eccDNA and remain similarly obscure. We therefore look on the present results as a stimulus for future work on eccDNA chromatin organization and epigenetics, investigations of transcription of eccDNAs of different sizes and superhelix densities and the possibility for subsequent translation into peptides or proteins with biological function.

Supplementary Data are available.

| System | Ions | Starting Structure | Simulation Length (ns) | Colour |
| --- | --- | --- | --- | --- |
| Dinucleosome | $\text{Ca}^{2+}$ | After 1ns implicit MD | 100 | Black |
| Dinucleosome | $\text{Na}^+$ | End of dinucleosome in $\text{Ca}^{2+}$ | 100 | Orange |
| Mononucleosome | $\text{Ca}^{2+}$ | After 1ns implicit MD | 100 | Purple |
| Naked microDNA | $\text{Na}^+$ | After 1.5ns implicit MD | 175 | Yellow |
| Naked microDNA | $\text{Ca}^{2+}$ | After 1.5ns implicit MD | 150 | Blue |
| Naked microDNA | $\text{Ca}^{2+}$ | After 2ns implicit MD | 175 | Red |
| Naked microDNA | $\text{Na}^+$ | End of naked eccDNA in $\text{Ca}^{2+}$ | 150 | Green |

## DATA AVAILABILITY

Python scripts used for analysing the MD simulations and generating the plots are available at the GitHub repository: https://github.com/Victor-93/DNA_Minicircles_analysis.

## SUPPLEMENTARY INFORMATION

### Model Building

The “figure 8” dinucleosome conformation was generated by 1) aligning Nuc A to Nuc B , 2) rotating Nuc B 180° about the axis joining the centers of the first and last base pairs, and 3) rotating Nuc B 180° about the axis joining the centers of mass of the histone core of Nuc A and Nuc B. A small translation was also introduced along the axis connecting the centers of mass of the histone cores so that the terminal base pairs of Nuc A and Nuc B were not overlapping. This “flip & twist” procedure provided a structure that contained the desired sequence, two nucleosomes each conforming to the 1KX5 x-ray structure including tails, arranged in a figure 8 structure with relaxed linker DNA.

The tails on the two nucleosomes in the dinucleosome-micro-eccDNA structure were modified by methylation of K9 and K27. Parameters from the AMBER ff14SB and ff14SB_modAA forcefields were used to describe standard and modified amino acids within the nucleosomes, and the AMBER BSC1 forcefield was used for the DNA. The system was solvated by TIP3P water.

## Supplementary Movies

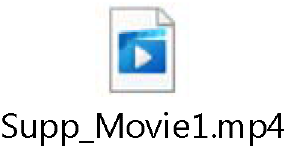

**Supplementary Movie 1:** Implicit solvent MD simulation showing the relaxation of the microDNA when one nucleosome is removed.

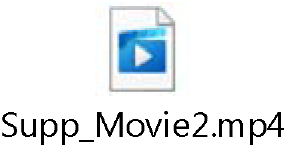

**Supplementary Movie 2:** Implicit solvent MD simulation showing the relaxation of the microDNA when two nucleosomes are removed.

## Supplementary Figures

**Supplementary Figure 1:**
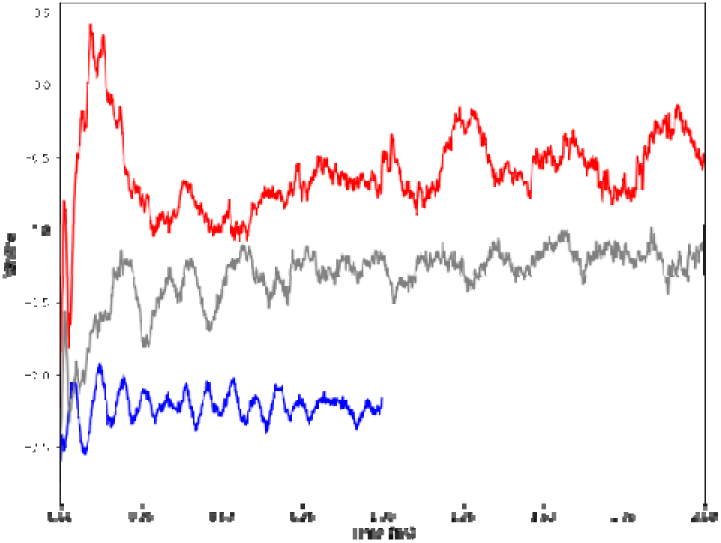
Time dependence of the writhe in implicitly solvated MD: Dinucleosome micro-eccDNA is shown in blue, mononucleosome micro-eccDNA in grey and naked micro-eccDNA in red.

**Supplementary Figure 2:**
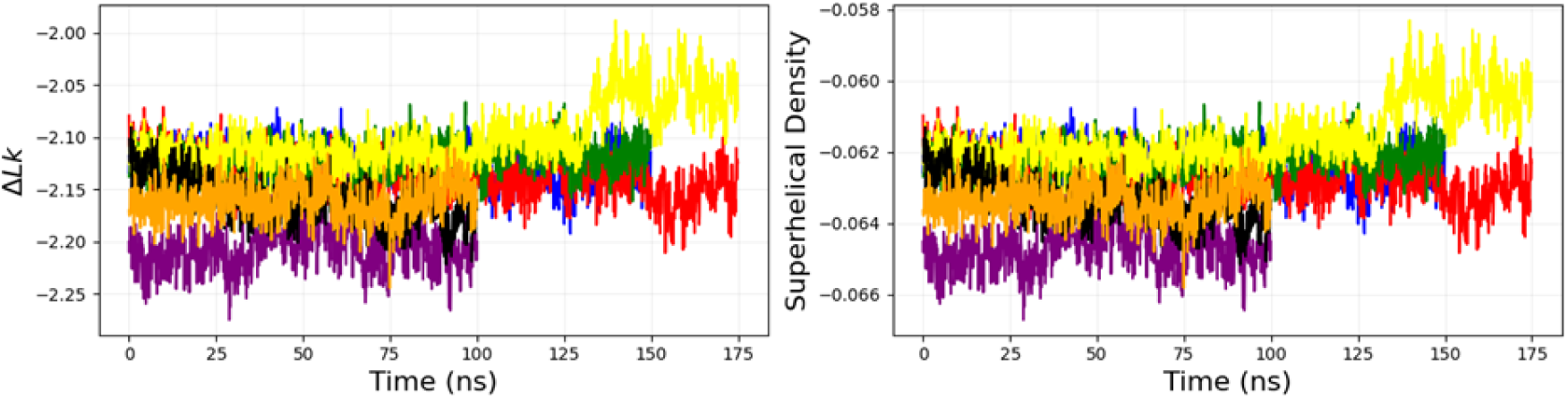
Time dependence of the linking number and superhelical densities for dinucleosome in CaCl_2_(black) and NaCl (orange); mononucleosome in CaCl_2_ (purple); and naked micro-eccDNA in CaCl_2_ (blue/red) and in NaCl (yellow/green.

**Supplementary Figure 3:**
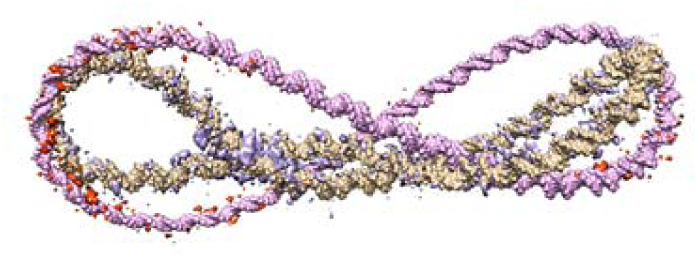
Superposition of structures and positive ion density maps for Ca^2+^ and Na^+^, superimposed on the average structure under CaCl_2_ (orange) and NaCl (purple). (density of Ca^2+^ shown in violet and Na^+^ density shown in red).

**Supplementary Figure 4:**
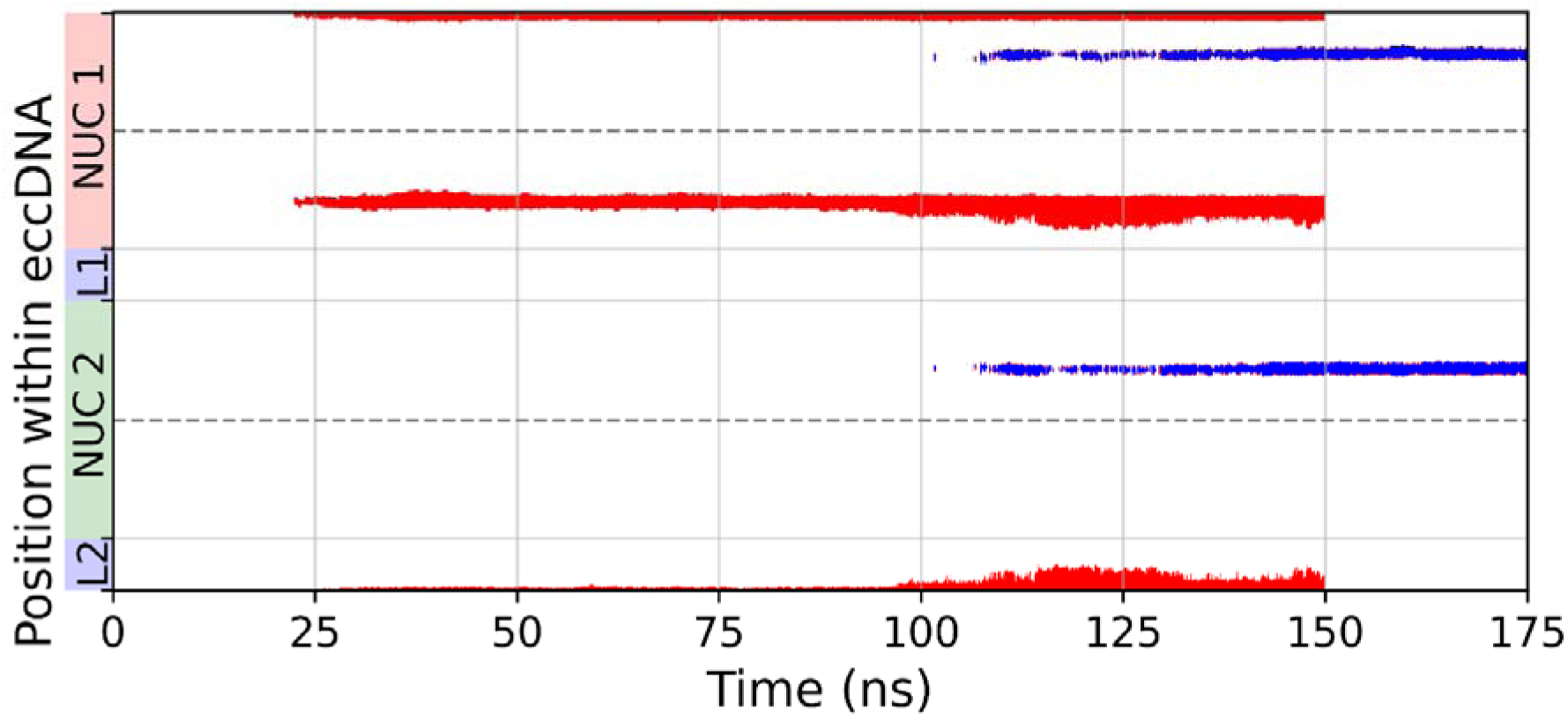
Close crossings in divalent ions for naked microDNA. Coloured points (blue and red points from two different replicas) indicate base pairs within 10 Å of base pairs that are more than 90 bases away, calculated from the centre line obtained using the wrLine program.

**Supplementary Figure S5:**
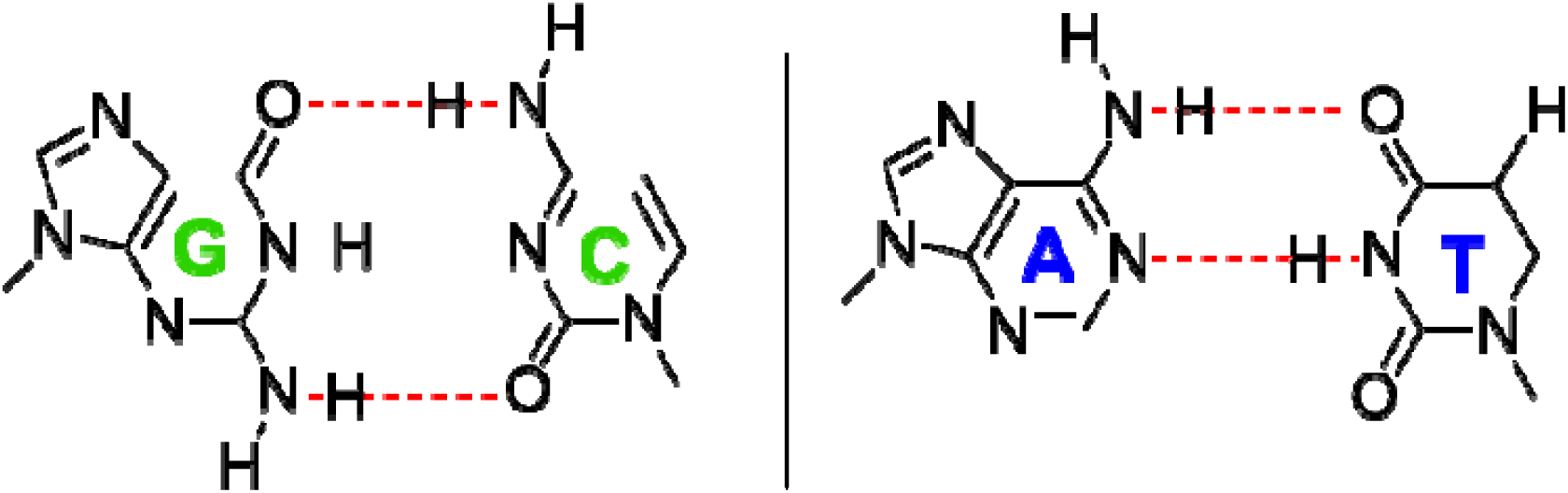
Definition of hydrogen bond distances. Hydrogen bond distances are calculated as the average distance between heavy atoms indicated by the red dashed line. Distances larger than 4Å are classified as DNA defects.

## REFERENCES

1. Shoura, M.J., Gabdank, I., Hansen, L., Merker, J., Gotlib, J., Levene, S.D. and Fire, A.Z. (2017) Intricate and Cell Type-Specific Populations of Endogenous Circular DNA (eccDNA) in Caenorhabditis elegans and Homo sapiens. G3 (Bethesda), 7, 3295–3303.

2. Paulsen, T., Malapati, P., Shibata, Y., Wilson, B., Eki, R., Benamar, M., Abbas, T. and Dutta, A. (2021) MicroDNA levels are dependent on MMEJ, repressed by c-NHEJ pathway, and stimulated by DNA damage. Nucleic Acids Res, 49, 11787–11799.

3. Bailey, C., Shoura, M.J., Mischel, P.S. and Swanton, C. (2020) Extrachromosomal DNA- relieving heredity constraints, accelerating tumour evolution. Ann Oncol, 31, 884–893.

4. Carroll, S.M., DeRose, M.L., Gaudray, P., Moore, C.M., Needham-Vandevanter, D.R., Von Hoff, D.D. and Wahl, G.M. (1988) Double minute chromosomes can be produced from precursors derived from a chromosomal deletion. Mol Cell Biol, 8, 1525–1533.

5. Benner, S.E., Wahl, G.M. and Von Hoff, D.D. (1991) Double minute chromosomes and homogeneously staining regions in tumors taken directly from patients versus in human tumor cell lines. Anticancer Drugs, 2, 11–25.

6. Cohen, S., Agmon, N., Sobol, O. and Segal, D. (2010) Extrachromosomal circles of satellite repeats and 5S ribosomal DNA in human cells. Mob DNA, 1, 11.

7. Cohen, S., Houben, A. and Segal, D. (2008) Extrachromosomal circular DNA derived from tandemly repeated genomic sequences in plants. Plant J, 53, 1027–1034.

8. Cohen, S., Regev, A. and Lavi, S. (1997) Small polydispersed circular DNA (spcDNA) in human cells: association with genomic instability. Oncogene, 14, 977–985.

9. Cohen, S. and Segal, D. (2009) Extrachromosomal circular DNA in eukaryotes: possible involvement in the plasticity of tandem repeats. Cytogenet Genome Res, 124, 327–338.

10. Cohen, S., Yacobi, K. and Segal, D. (2003) Extrachromosomal circular DNA of tandemly repeated genomic sequences in Drosophila. Genome Res, 13, 1133–1145.

11. Gaubatz, J.W. and Flores, S.C. (1990) Tissue-specific and age-related variations in repetitive sequences of mouse extrachromosomal circular DNAs. Mutat Res, 237, 29–36.

12. Kapitonov, V.V. and Jurka, J. (2007) Helitrons on a roll: eukaryotic rolling-circle transposons. Trends Genet, 23, 521–529.

13. Kinoshita, Y., Ohnishi, N., Yamada, Y., Kunisada, T. and Yamagishi, H. (1985) Extrachromosomal circular DNA from nuclear fraction of higher plants. Plant and cell physiology, 26, 1401–1409.

14. Morton, A.R., Dogan-Artun, N., Faber, Z.J., MacLeod, G., Bartels, C.F., Piazza, M.S., Allan, K.C., Mack, S.C., Wang, X., Gimple, R.C. et al. (2019) Functional Enhancers Shape Extrachromosomal Oncogene Amplifications. Cell, 179, 1330–1341 e1313.

15. Cohen, S. and Méchali, M. (2002) Formation of extrachromosomal circles from telomeric DNA in Xenopus laevis. EMBO Rep, 3, 1168–1174.

16. Demeke, M.M., Foulquie-Moreno, M.R., Dumortier, F. and Thevelein, J.M. (2015) Rapid evolution of recombinant Saccharomyces cerevisiae for Xylose fermentation through formation of extra-chromosomal circular DNA. PLoS Genet, 11, e1005010.

17. Paulsen, T., Shibata, Y., Kumar, P., Dillon, L. and Dutta, A. (2019) Small extrachromosomal circular DNAs, microDNA, produce short regulatory RNAs that suppress gene expression independent of canonical promoters. Nucleic Acids Res, 47, 4586–4596.

18. Azam, S., Yang, F. and Wu, X. (2025) Finding functional microproteins. Trends Genet, 41, 107–118.

19. Apcher, S., Tovar-Fernadez, M., Ducellier, S., Thermou, A., Nascimento, M., Sroka, E. and Fahraeus, R. (2022) mRNA translation from an antigen presentation perspective: A tribute to the works of Nilabh Shastri. Mol Immunol, 141, 305–308.

20. De Lucia, F., Alilat, M., Sivolob, A. and Prunell, A. (1999) Nucleosome dynamics. III. Histone tail-dependent fluctuation of nucleosomes between open and closed DNA conformations. Implications for chromatin dynamics and the linking number paradox. A relaxation study of mononucleosomes on DNA minicircles. J Mol Biol, 285, 1101–1119.

21. Prunell, A., Alilat, M. and De Lucia, F. (1999) Nucleosome structure and dynamics. The DNA minicircle approach. Methods Mol Biol, 119, 79–101.

22. Irobalieva, R.N., Fogg, J.M., Catanese, D.J., Jr., Sutthibutpong, T., Chen, M., Barker, A.K., Ludtke, S.J., Harris, S.A., Schmid, M.F., Chiu, W. et al. (2015) Structural diversity of supercoiled DNA. Nat Commun, 6, 8440.

23. Pyne, A.L.B., Noy, A., Main, K.H.S., Velasco-Berrelleza, V., Piperakis, M.M., Mitchenall, L.A., Cugliandolo, F.M., Beton, J.G., Stevenson, C.E.M., Hoogenboom, B.W. et al. (2021) Base-pair resolution analysis of the effect of supercoiling on DNA flexibility and major groove recognition by triplex-forming oligonucleotides. Nat Commun, 12, 1053.

24. Cordero, P., Parikh, V.N., Chin, E.T., Erbilgin, A., Gloudemans, M.J., Shang, C., Huang, Y., Chang, A.C., Smith, K.S., Dewey, F. et al. (2019) Pathologic gene network rewiring implicates PPP1R3A as a central regulator in pressure overload heart failure. Nat Commun, 10, 2760.

25. Stolz, R.C. and Bishop, T.C. (2010) ICM Web: the interactive chromatin modeling web server. Nucleic Acids Res, 38, W254–261.

26. Davey, C.A., Sargent, D.F., Luger, K., Maeder, A.W. and Richmond, T.J. (2002) Solvent mediated interactions in the structure of the nucleosome core particle at 1.9 a resolution. J Mol Biol, 319, 1097–1113.

27. D.A. Case, K.B., I.Y. Ben-Shalom, S.R. Brozell, D.S. Cerutti, T.E. Cheatham, III, V.W.D. Cruzeiro, T.A. Darden, R.E. Duke, G. Giambasu, M.K. Gilson, H. Gohlke, A.W. Goetz, R Harris, S. Izadi, S.A. Izmailov, K. Kasavajhala, A. Kovalenko, R. Krasny, T. Kurtzman, T.S. Lee, S. LeGrand, P. Li, C. Lin, J. Liu, T. Luchko, R. Luo, V. Man, K.M. Merz, Y. Miao, O. Mikhailovskii, G. Monard, H. Nguyen, A. Onufriev, F. Pan, S. Pantano, R. Qi, D.R. Roe, A. Roitberg, C. Sagui, S. Schott-Verdugo, J. Shen, C.L. Simmerling, N.R. Skrynnikov, J. Smith, J. Swails, R.C. Walker, J. Wang, L. Wilson, R.M. Wolf, X. Wu, Y. Xiong, Y. Xue, D.M. York and P.A. Kollman (2020) AMBER 2020

28. Case, D.A., Aktulga, H.M., Belfon, K., Cerutti, D.S., Cisneros, G.A., Cruzeiro, V.W.D., Forouzesh, N., Giese, T.J., Götz, A.W., Gohlke, H. et al. (2023) AmberTools. Journal of Chemical Information and Modeling, 63, 6183–6191.

29. Sutthibutpong, T., Noy, A. and Harris, S. (2016) Atomistic Molecular Dynamics Simulations of DNA Minicircle Topoisomers: A Practical Guide to Setup, Performance, and Analysis. Methods Mol Biol, 1431, 195–219.

30. Sutthibutpong, T., Harris, S.A. and Noy, A. (2015) Comparison of molecular contours for measuring writhe in atomistic supercoiled DNA. J Chem Theory Comput, 11, 2768–2775.

31. Gowers, R., Linke, M., Barnoud, J., Reddy, T., Melo, M., Seyler, S., Domański, J., Dotson, D., Buchoux, S., Kenney, I. et al. (2016), Proceedings of the 15th Python in Science Conference, pp. 98–105.

32. Michaud-Agrawal, N., Denning, E.J., Woolf, T.B. and Beckstein, O. (2011) MDAnalysis: a toolkit for the analysis of molecular dynamics simulations. J Comput Chem, 32, 2319–2327.

33. Dorman, C.J. (2006) Dna Supercoiling and Bacterial Gene Expression. Science Progress, 89, 151–166.

34. Bliska, J.B. and Cozzarelli, N.R. (1987) Use of site-specific recombination as a probe of DNA structure and metabolism in vivo. J Mol Biol, 194, 205–218.

35. Fogg, J.M., Judge, A.K., Stricker, E., Chan, H.L. and Zechiedrich, L. (2021) Supercoiling and looping promote DNA base accessibility and coordination among distant sites. Nature Communications, 12, 5683.

36. Matek, C., Ouldridge, T.E., Doye, J.P.K. and Louis, A.A. (2015) Plectoneme tip bubbles: Coupled denaturation and writhing in supercoiled DNA. Scientific Reports, 5, 7655.

37. Sutthibutpong, T., Matek, C., Benham, C., Slade, G.G., Noy, A., Laughton, C., K. Doye, J.P., Louis, A.A. and Harris, S.A. (2016) Long-range correlations in the mechanics of small DNA circles under topological stress revealed by multi-scale simulation. Nucleic Acids Research, 44, 9121–9130.

38. Galindo-Murillo, R., Roe, D.R. and Cheatham, T.E., 3rd. (2014) On the absence of intrahelical DNA dynamics on the mus to ms timescale. Nat Commun, 5, 5152.

39. Várnai, P. and Timsit, Y. (2010) Differential stability of DNA crossovers in solution mediated by divalent cations. Nucleic Acids Research, 38, 4163–4172.

40. Timsit, Y. and Várnai, P. (2010) Helical chirality: a link between local interactions and global topology in DNA. PLoS One, 5, e9326.

41. Bednar, J., Furrer, P., Stasiak, A., Dubochet, J., Egelman, E.H. and Bates, A.D. (1994) The twist, writhe and overall shape of supercoiled DNA change during counterion-induced transition from a loosely to a tightly interwound superhelix. Possible implications for DNA structure in vivo. J Mol Biol, 235, 825–847.

42. Sottas, P.E., Larquet, E., Stasiak, A. and Dubochet, J. (1999) Brownian dynamics simulation of DNA condensation. Biophys J, 77, 1858–1870.

43. Bowman, G.D. and Poirier, M.G. (2015) Post-Translational Modifications of Histones That Influence Nucleosome Dynamics. Chemical Reviews, 115, 2274–2295.

